# Assessing student success in a Peer Assisted Learning program using propensity score matching

**DOI:** 10.1101/2021.06.07.447443

**Authors:** Corey Shanbrom, Michelle Norris, Caitlin Esgana, Matthew Krauel, Vincent Pigno, Jennifer Lundmark

## Abstract

The Peer Assisted Learning program at Sacramento State University (PAL) was established in 2012 with one section supporting introductory chemistry. It now serves 17 gatekeeper courses in Biology, Chemistry, Mathematics, Physics, and Statistics, enrolling approximately 1,400 students annually. Adapting the Peer-Led Team Learning model, PAL Facilitators do not teach, tutor, or even confirm answers; they do ask scaffolding questions, provide encouragement, and ensure that all group members participate in problem-solving. Here we assess the efficacy of the program in terms of student success in the parent course. As PAL is an opt-in program, we employ propensity score matching techniques to account for confounding factors. Our analysis of 11 classes shows that PAL provides an average course GPA bump ranging from .23 to .71 grade points (mean .42). Compared to the non-PAL baseline course GPA, this amounts to an increase of 9% to 51% (mean 23%). We consider data from over 25,000 students, and our propensity score analysis uses over 10,000 students (4,519 PAL, 5,814 non-PAL) for whom appropriate matches could be found.

## Introduction

The positive impact of Peer-Led Team Learning (PLTL) on student success in STEM courses is well established in the literature (Arendale, 2019). However, while enrollment is mandatory in the original PLTL model (Gosser et al., 2001), in many institutions students opt-in to PLTL programs (Frey et al., 2018), creating a number of possible confounding variables. Put simply, it may be that the type of students who choose to participate were already more likely to earn higher grades or pass at higher rates than their peers who did not voluntarily join the program; therefore one cannot conclude that the program is itself responsible for their success. This concern has been recognized by a number of researchers (Frey et al., 2018; Chan & Bauer, 2015). The statistical method of propensity score matching can help solve this problem by pairing groups of students who choose to participate in the intervention with groups who do not participate, but who differ in no known other academic or demographic ways. Here we apply propensity score matching analysis to such a voluntary PLTL program at a large public university in order to tease out the true efficacy of the program itself.

California State University, Sacramento, also known as Sacramento State, is one of 23 campuses in the California State University system. Sacramento State is a primarily undergraduate institution, enrolling over 30,000 undergraduate students annually. It is officially recognized as an Asian American Native American Pacific Islander Serving Institution (AANAPISI) and a Hispanic Serving Institution (HSI), and was ranked the 2^nd^ most diverse regional university in the West (U.S. News & World Report, 2021). Approximately 32% of students are first in their family to attend college.

The Peer Assisted Learning program at Sacramento State (PAL) was launched in 2012 to offer academic support to students in gatekeeper science and mathematics courses. It serves approximately 1,400 students per year in 17 such courses. The 11 courses analyzed in this study appear in Table 1. The program also serves college algebra (MATH 12), introductory statistics (STAT 1), calculus 3 (MATH 32), advanced organic chemistry (CHEM 124), physiology (BIO 25), and general physics (PHYS 5A), but these are newer additions and there are not yet enough data to conduct meaningful propensity score analysis. The PAL program is housed within the Center for Science and Math Success with several related programs within the College of Natural Sciences and Mathematics. An affiliate program is currently being developed in the College of Engineering and Computer Science.

**Table 1:** PAL course DFW rates, Spring 2016-Fall 2019 (8 semesters).

| Course code | Content | DFW rate |
| --- | --- | --- |
| BIO 22 | Anatomy | 35% |
| BIO 121 | Molecular Cell Biology | 24% |
| BIO 131 | Physiology | 21% |
| BIO 184 | Genetics | 13% |
| CHEM 1A | General Chemistry 1 | 40% |
| CHEM 1B | General Chemistry 2 | 32% |
| CHEM 4 | Chemical Calculations | 35% |
| CHEM 24 | Organic Chemistry | 37% |
| MATH 29 | Precalculus | 25% |
| MATH 30 | Calculus 1 | 26% |
| MATH 31 | Calculus 2 | 29% |

Note that there are a number of “Peer Assisted Learning” programs across the country (Arendale, 2019), with the first founded at University of Minnesota in 2006 (Arendale, 2014), but in this paper we use the acronym “PAL” exclusively to refer to the Sacramento State program for readability.

While PAL is based on the PLTL model (Lundmark et al., 2017), a number of features distinguish it from similar programs at other institutions, including those also known as “Peer Assisted Learning”. Following PLTL, students work in small groups (3-4 students) on worksheets written by Sacramento State course faculty. Groups work on whiteboards with one pen which rotates among group members. The peer Facilitator ensures that the pen cycles regularly and all group members are participating in the problem solving. The Facilitator also offers encouragement and positive reinforcement. Facilitators do not, however, teach, tutor, or even confirm whether answers are correct. Instead, they can ask scaffolding questions to help guide students toward solving the problems on their own.

A typical PAL section consists of one Facilitator and approximately 15 students. Each section meets for 50 minutes twice per week and runs as an independent 1-unit class graded Credit/No Credit. Each section is directly connected to a primary math or science course. Importantly for our statistical analysis, enrollment in PAL is completely voluntary. However, once enrolled, attendance is mandatory. The program marketing is that PAL is for everyone; it is not remediation.

Each of the approximately 70 PAL Facilitators also holds regular office hours and attends lectures for the parent course. They can earn extra paid hours by holding review sessions, attending relevant professional development or trainings (Dreamer Ally, Safe Zone, etc.), or by participating in leadership (e.g., Cultural Competency Ambassadors). Office hours and review sessions are open to all students from the parent course, regardless of enrollment in a PAL section.

The program also has a unique leadership structure. PAL is led by a team of four faculty members, assisted by one part-time staff member. But much of the day-to-day is run by a team of 3-4 Supervisory Facilitators, who are experienced Facilitators that show exceptional professionalism and leadership potential. They serve as faculty liaisons, organize and conduct all classroom observations, manage the logistics of office hours, and host social events and parties (PALidays). Moreover, each of the 17 courses possesses a Lead Facilitator in charge of student recruitment, communication with course faculty, and leading the weekly run-through of the upcoming worksheets with an emphasis on practicing scaffolding.

A unique aspect of the PAL program among various supplemental instruction models is the requirement that all Facilitators conduct action research projects. All Facilitators take an upper division 2-unit course, Honors Seminar in Peer Learning, which meets Wednesday nights for two hours with all four faculty. In addition to the worksheet run-through, this seminar includes guest speakers, PAL panels, and ongoing pedagogical trainings (self-efficacy, growth-mindset, cultural competency, metacognition, etc.). However, the seminar primarily offers Facilitators an opportunity to conduct education research projects within their PAL section. Interdisciplinary teams of approximately five Facilitators use backward design to plan these projects; they are asked to start by identifying what they really want PAL students to be able to do or understand, then decide how they will determine if they’ve reached these goals, then design learning experiences accordingly (Wiggins & McTighe, 2005). The project methodologies and backgrounds are developed in Fall. In Spring, the designed interventions are implemented, data is collected and analyzed, and posters are presented in a session open to the campus community. Many of these projects have led to structural changes within the program as Facilitators discover what seems to work best in their own classrooms. Some of these projects have led to grants, conference presentations, and even scholarly publications (Lundmark et al., 2017).

The continual improvement of the PAL program, thanks to Facilitator research projects, has been crucial in institutionalizing the program on campus and securing funding, and allows the program to attract excellent student leaders. Raw data show that students who opt-in to the program outperform their peers who do not. However, determining whether this is a true “PAL effect”, rather than the result of stronger students opting in, requires advanced statistical techniques detailed in the next section.

## Methods

A propensity score analysis was conducted using the Matching (Sekhon, 2011) and cobalt (Greifer, 2021) packages in R (R Core Team, 2019) to assess the effect of PAL supplemental instruction on course grades in 11 STEM courses (see Table 1). Propensity score adjustment was necessary since the data are observational and the characteristics of students who voluntarily enroll in PAL may differ in ways that may, independently of PAL, impact course grade compared to students who do not enroll in PAL. In propensity score analysis, variables related to both likelihood of PAL enrollment and course grade (confounders) are used in a logistic regression model to obtain a propensity score, which is a student’s likelihood of enrolling in PAL. Details on implementing propensity score matching techniques using R can be found in Leite (2016). Such methods have been used to evaluate a number of STEM student success programs: non-peer-led interventions in an introductory physics course (Rose, 2013), a PLTL program in STEM courses in urban high schools (Thomas et al., 2015), non-peer-led college math readiness interventions (Hodara, 2013), and STEM retention (Windsor et al., 2015). Moreover, Carlsen et al. (2016) employed “matched-pairs design” to evaluate a PLTL program in which participants were matched with non-participants who were “taking the same unit, often with the same instructor.” To our knowledge, our study is the first application of propensity score matching to Peer Assisted Learning or Peer Led Team Learning programs at the postsecondary level.

We performed a customized propensity score matching analysis for each of the 11 courses. For each course, we began with 174 variables for every student enrolled in the parent course for relevant dates: since the first PAL section was offered (Spring 2012 for the oldest) until Spring 2019. This large set of variables was reduced in three ways. First, we analyzed missingness and handled missing data. We handled missing data by removing variables with insufficient data. Second, our subjective judgment was used to narrow the pool of remaining variables down to those likely to be confounders. We included all variables likely to be correlated with outcome even if it was uncertain whether they were related to likelihood of enrolling in PAL. This allowed for a more precise estimate of the PAL treatment effect. Both of these reductions were specific to the course. Then students that were missing a value from any of the remaining variables were removed. This resulted in a smaller dataset without missing values. Third, covariates were selected for the propensity model using stepwise regression. In some cases, subjective judgment was used to add new variables which were likely relevant, although not included in our original list of 174 variables. For example, in our analysis of Chem 1B, we added a variable for students’ grades in Chem 1A.

Using the propensity score model, all students in the dataset, PAL and non-PAL, were assigned a propensity score, representing their likelihood of enrolling in PAL. Then, each PAL student was matched to one or more non-PAL students who had similar propensity scores. When a PAL student had more than one suitable match among the non-PAL students, all non-PAL students were taken as matches and weighted appropriately in the final estimated PAL effect. After matching, the PAL and matched non-PAL groups were compared to determine whether the distribution of each covariate was similar between the two groups. This is called a balance check. If the standardized difference between the non-PAL and PAL means is less than 0.10 then the strong criteria in (Leite, 2016) is met for covariate balance. If the standardized difference is under 0.25, then a more lenient criteria is met.

The difference in the average grade for the matched PAL and non-PAL data was then calculated. Courses without statistically reliable data (due to an insufficient number of PAL students) were excluded. The results are presented in the next section. Much more detailed descriptions of our methods, including some of our code and R packages used, can be found in the R Markdown files available online (Shanbrom, 2021). In addition to Leite (2016), a number of research papers were used in our analysis (Brookhart et al., 2006; Austin, 2011; Liu et al., 2013; Zhang et al., 2019).

## Results

Our main results appear in Table 2, where for each course we display the propensity score adjusted mean course GPA for both students who opted into the course PAL and the matched students who did not. A visualization of this data appears in Figure 1. The table shows that PAL provides an average course GPA bump ranging from .23 to .71 grade points (mean .42). Compared to the non-PAL baseline course GPA, this amounts to an increase of 9% to 51% (mean 23%). We conclude that PAL is effective in increasing student success in each of these courses, and that this effect is not confounded by the PAL students’ propensity to enroll in the program.

**Figure 1:**
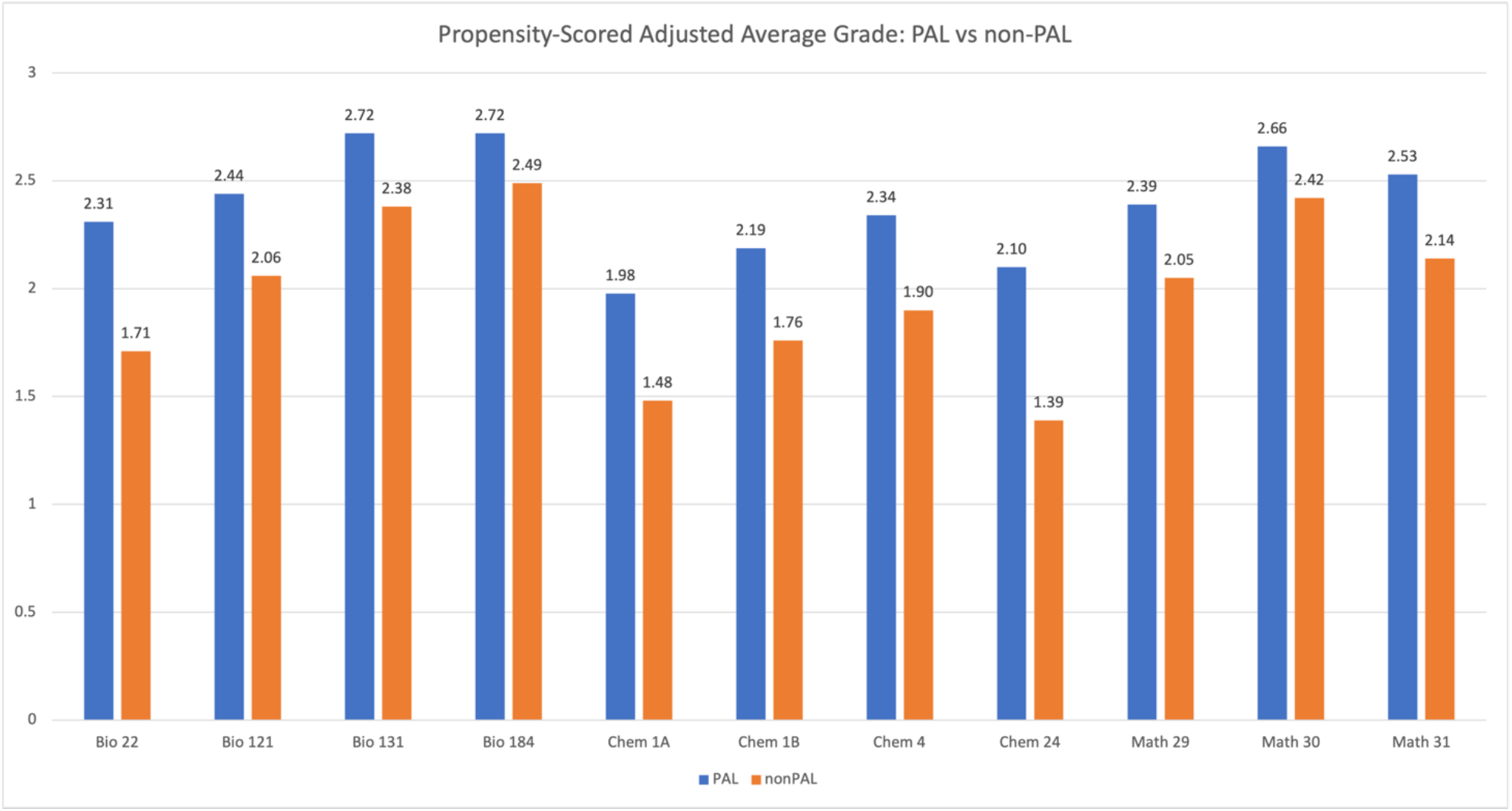
Visualization of the PAL bump after propensity score matching.

**Table 2:**
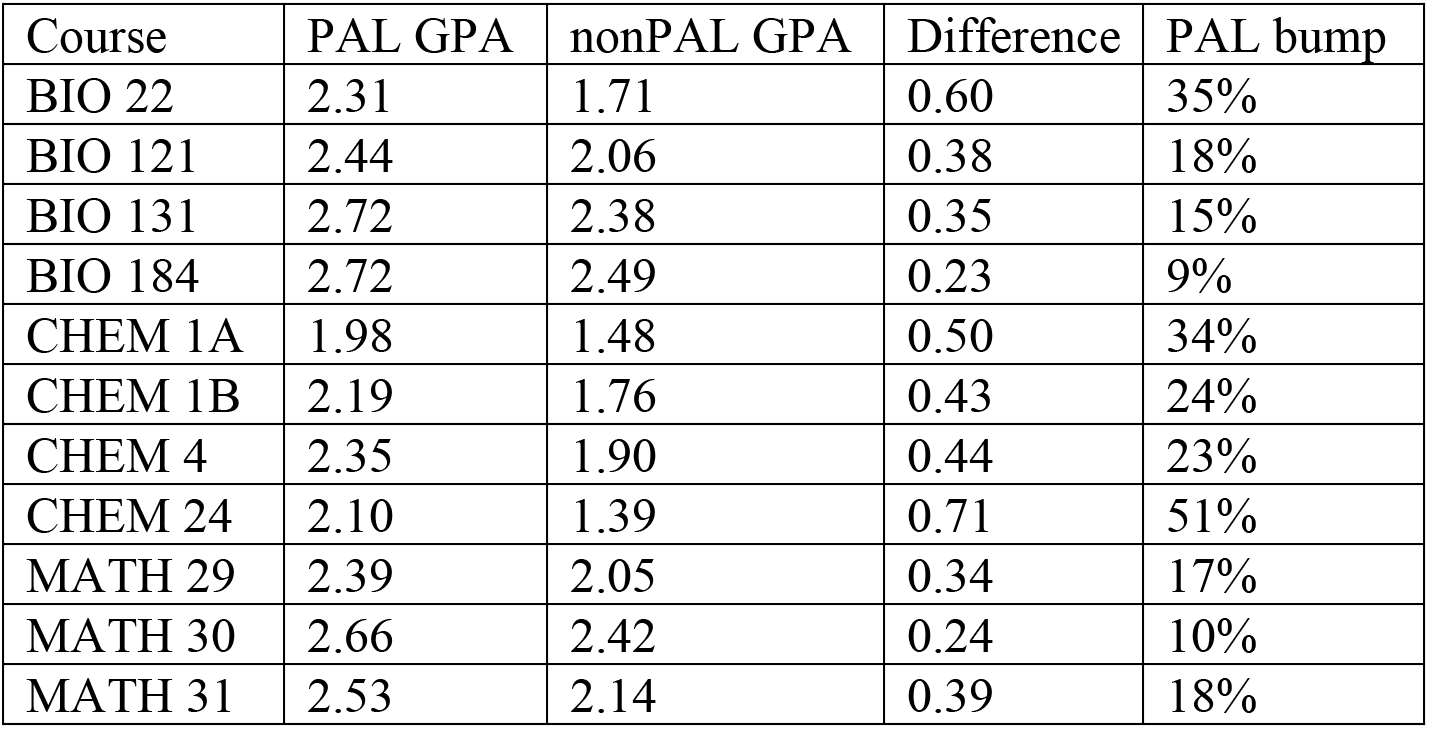
Main result: propensity score adjusted mean course GPA increase from PAL. These data are displayed visually in Figure 1. Course content appears in Table 1.

| Course | PAL GPA | nonPAL GPA | Difference | PAL bump |
| --- | --- | --- | --- | --- |
| BIO 22 | 2.31 | 1.71 | 0.60 | 35% |
| BIO 121 | 2.44 | 2.06 | 0.38 | 18% |
| BIO 131 | 2.72 | 2.38 | 0.35 | 15% |
| BIO 184 | 2.72 | 2.49 | 0.23 | 9% |
| CHEM 1A | 1.98 | 1.48 | 0.50 | 34% |
| CHEM 1B | 2.19 | 1.76 | 0.43 | 24% |
| CHEM 4 | 2.35 | 1.90 | 0.44 | 23% |
| CHEM 24 | 2.10 | 1.39 | 0.71 | 51% |
| MATH 29 | 2.39 | 2.05 | 0.34 | 17% |
| MATH 30 | 2.66 | 2.42 | 0.24 | 10% |
| MATH 31 | 2.53 | 2.14 | 0.39 | 18% |

Table 3 provides the important statistical context needed to interpret Table 2. Standard errors, p-values, sensitivity, and sample sizes (N) are provided for each of the 11 courses. The standard errors and p-values are small enough to conclude that our results are indeed statistically significant. The sensitivity column displays the number Γ which can be interpreted as follows. An unknown confounder which increases the odds of being in PAL by more than Γ is enough to change the treatment effect from significant to non-significant.

**Table 3:** Relevant statistics; N values after propensity score matching.

| Course | p-value | Standard error | Sensitivity | N(PAL) | N(nonPAL) |
| --- | --- | --- | --- | --- | --- |
| BIO 22 | 1.28e-6 | 0.13 | 1.6 | 326 | 332 |
| BIO 121 | 9.17e-5 | 0.10 | 1.4 | 322 | 370 |
| BIO 131 | 4.24e-7 | 0.07 | 1.5 | 469 | 751 |
| BIO 184 | 5.03e-2 | 0.14 | 1.1 | 143 | 153 |
| CHEM 1A | 2.43e-13 | 0.07 | 1.8 | 757 | 711 |
| CHEM 1B | 8.59e-9 | 0.08 | 1.8 | 573 | 457 |
| CHEM 4 | 2.19e-7 | 0.09 | 1.7 | 530 | 680 |
| CHEM 24 | 3.16e-3 | 0.26 | 1.8 | 126 | 50 |
| MATH 29 | 4.77e-6 | 0.08 | 1.4 | 475 | 850 |
| MATH 30 | 4.76e-4 | 0.07 | 1.3 | 506 | 870 |
| MATH 31 | 2.81e-5 | 0.10 | 1.5 | 292 | 590 |

Note that the sample sizes displayed in Table 3 represent the number of students actually used in the propensity score analysis. This is much smaller than the original raw sample size since we drop students with missing data as well as students from one pool (PAL or non-PAL) who do not closely match at least one student from the other pool. Moreover, students in one pool can be matched to multiple students in the other pool. For example, in Chem 1A, the original pool contained 1561 non-PAL students and 762 PAL students, but the propensity score matching analysis used subsets of 711 non-PAL students and 757 PAL students. These latter numbers are displayed in the table. Of the 757 PAL students, 337 were matched one-to-one with non-PAL students and 420 were matched one-to-many with non-PAL students. This is displayed visually by the histogram in Figure 2.

**Figure 2:**
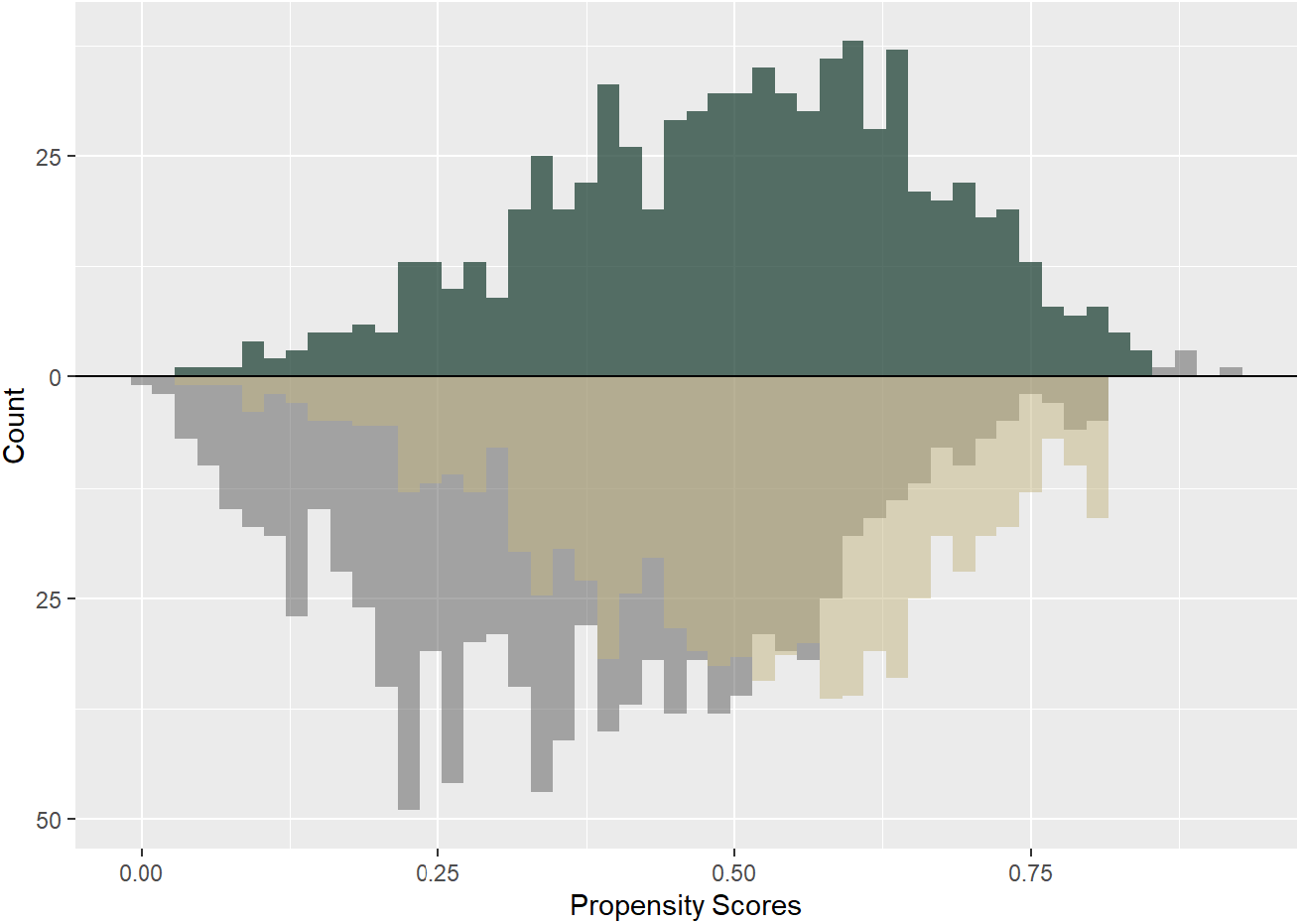
Visualization of propensity score matching in Chem 1A. Top shows PAL students, bottom shows non-PAL students. Grey represents the original pools before matching. Matched PAL students are colored green; matched non-PAL students are colored gold. Due to the one-to-many matching, matched non-PAL students are counted with multiplicity: dark gold is the overlap of gold with grey.

It is important to note that the math classes analyzed here represent a mixture of two treatments. In Math 29, 30, and 31, certain PAL classes were associated to special lecture courses called Learning Communities, which were composed entirely of first-year students and for which PAL enrollment was mandatory. While the Learning Communities themselves were opt-in, the overall course experience was different enough for these students that we consider their PAL participation as a different kind of treatment. Excluding the Learning Communities would significantly shrink the PAL student pool. For example, in Math 30, including the Learning Community students yields 508 matched PAL students, as shown in Table 3, while excluding the Learning Communities yields only 195 matched PAL students. For this reason, we chose to include the Learning Communities and acknowledge that the analysis for these math courses holds for a mix of two treatments.

Table 4 shows the 13 variables that were selected as covariates in our analysis of Chem 1B. Some of these covariates are categorical, some numerical, and some binary. They are listed by decreasing standard mean difference (SMD) between the unmatched PAL versus non-PAL populations, where the SMD of a numerical covariate is defined as the difference in the PAL and the non-PAL means of the covariate divided by the standard deviation of the covariate. Figure 3 shows the effect of propensity score matching on reducing the SMDs. Note that the SMDs are smaller for the matched populations for nearly all variables. Some majors or ethnicities had sparse categories that needed to be collapsed or grouped together to avoid complete separation in logistic regression. The list of covariates used and their corresponding SMDs depend on the course being analyzed. Additional detailed results for each class can be found in the R Markdown files available online (Shanbrom 2021).

**Figure 3:**
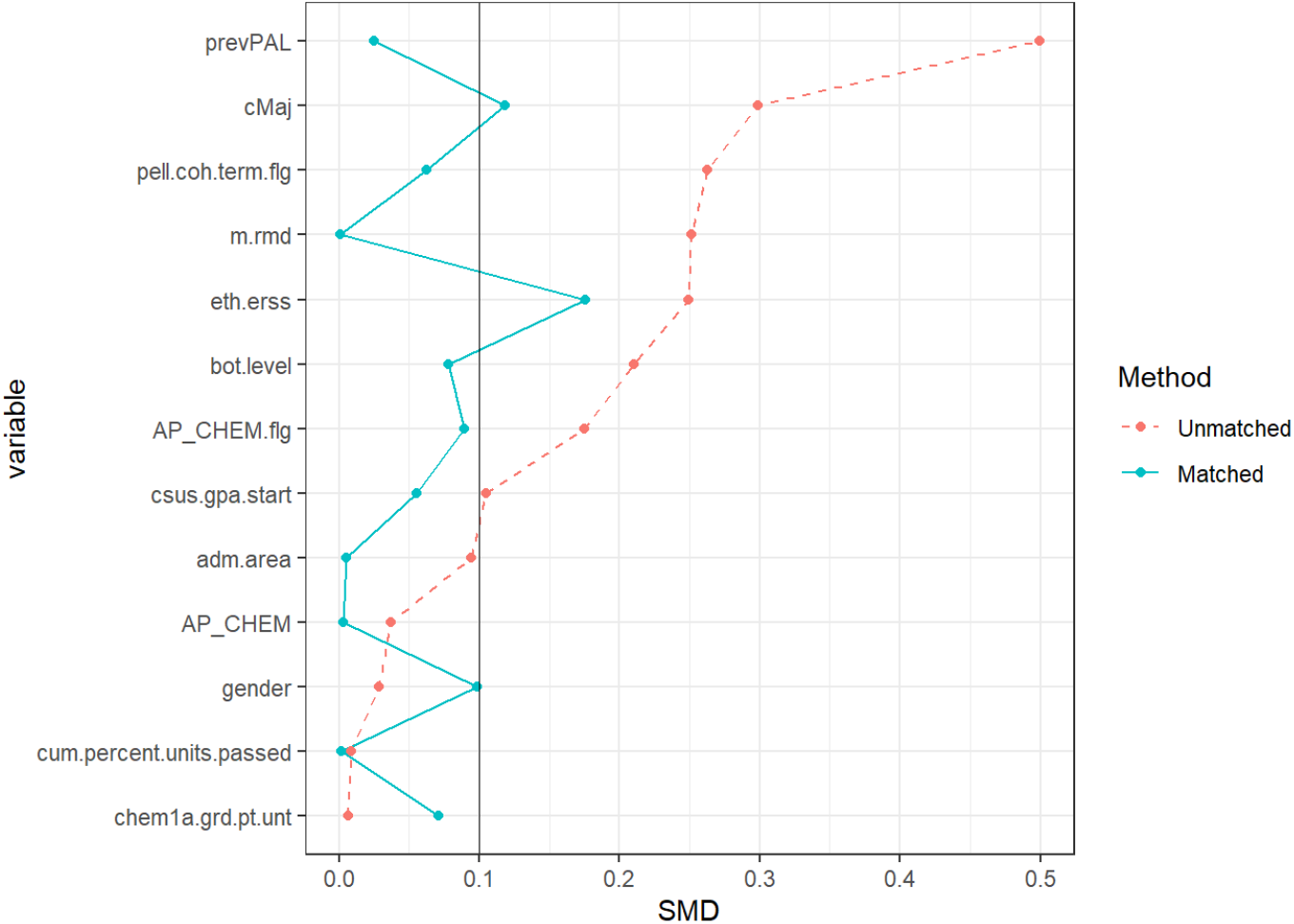
The 13 variables used in Chem 1B analysis; descriptions in Table 3. Plot maps the difference between the matched and unmatched PAL vs. non-PAL populations based on their respective SMDs and shows the reduced SMDs under matching in nearly all variables.

**Table 4:** Covariates used in Chem 1B analysis as provided by institutional data.

| Name | Description |
| --- | --- |
| prevPAL | Did student take PAL in at least one previous term |
| cMaj | Student's major as of census date |
| pell.coh.term.flg | Pell eligibility when student entered Sacramento State |
| m.rmd | Remedial in math |
| eth.erss | Ethnicity |
| bot.level | Class level at beginning of term |
| AP_CHEM.flg | Did student take AP Chemistry exam |
| csus.gpa.start | Sacramento State GPA at start of term |
| adm.area | Is student from Sacramento State local admission area |
| AP_CHEM | AP Chemistry exam score |
| gender | Gender |
| cum.percent.units.passed | Cumulative units passed/cumulative units taken |
| chem1a.grd.pt.unt | Course GPA in Chem 1A |

Finally, our analysis also provided participation data. We found that, in the classes and terms analyzed, a total of 7,180 students participated in PAL while 18,449 did not; thus the overall participation rate is approximately 28%. Of the 113 semester-courses included in the study, these rates varied considerably, from a low of 9% (Math 31, Spring 2016) to a high of 55% (Chem 24, Spring 2019).

In total, this analysis considered data from over 25,000 students. However, most of these students did not match well with students from the opposite pool or had too many missing values to be included in the analysis. The propensity score analysis itself used over 10,000 students (4,519 PAL, 5,814 non-PAL).

## Conclusion

We successfully conducted a propensity score analysis to assess the effect of PAL participation on course grades in 11 STEM courses. Our results show that students who choose to enroll in the PAL program earn significantly higher course grades than students with comparable social and academic backgrounds who do not enroll. Nevertheless, there are a number of shortcomings to this analysis and challenges to its replication or extension.

Most importantly, our propensity scoring was limited to the covariates available in our institutional database. While this includes a large number of variables (174), not all were meaningful, and there are obviously many other variables which could affect both a student’s course grade and their likelihood of enrolling in PAL. Such variables are known as unknown confounders. For example, our dataset does not contain information on the number of hours a student works per week, which may significantly affect the student’s ability or willingness to participate in PAL. The sensitivity displayed in Table 3 attempts to quantify the amount by which our results could depend on such unknown confounders (Rosenbaum, 2002).

For example, in Bio 121, the sensitivity analysis indicates that an unknown confounder which increases the odds of being in PAL by more than 1.4 is enough to change the treatment effect from significant to non-significant. Inspection of the covariates in the estimated propensity model for Bio 121 indicates that if there is an unknown confounder that has an effect on the propensity score similar to the effect of ethnicity, major or instructor observed in this analysis, the PAL effect would become non-significant. Thus, this finding is moderately sensitive to unknown confounders. Larger sensitivity values correspond to less sensitive results, so courses like Chem 1B and Chem 24 are less likely to have our results nullified by unknown confounders. On the other hand, Bio 184 is quite sensitive, and different choices of variables could potentially yield significantly different results.

Additionally, depending on the course, a number of important variables were removed due to large amounts of missingness. In Bio 121, for example, 46% of SAT scores and 41% of high school GPA values were missing. This may be tied to transfer status: a large number of students on our campus are transfer students who do not need to provide pre-secondary information, as their performance at their community college is taken as evidence of their preparedness. Since all students who had missing information on any included covariate were eliminated from the analysis, a balance had to be struck between retaining a sufficiently large pool of PAL and non-PAL students and retaining a sufficient number of important covariates. For Bio 121, grades in the prerequisite course, Bio 184, were available for most students and could reasonably be expected to provide a stronger measure of preparedness for Bio 121 than SAT scores. Thus, the removal of SAT scores from the propensity model was of little concern for Bio 121. However, for other courses, where the only available measure of preparedness for the course was SAT scores, the necessity of removing SAT scores due to missingness was of greater concern. In general, the hardest part of this analysis was accounting for the large amount of missing data. This presents challenges for extending or updating this work, as well as for other practitioners working with data sets even less robust than our own.

## Acknowledgements

This analysis was supported by NSF DUE 1644273, Sacramento State Academic Affairs, and Sacramento State Student Success Initiatives. We are also grateful to Joel Schwartz for proposing propensity analysis, and for his help wrangling and understanding the dataset on which this work is based.

## Bibliography

Arendale, D. (2014). Understanding the Peer Assistance Learning model: Student study groups in challenging college courses. International Journal of Higher Education, 3(2), 1–12.

Arendale, D. (2019). Postsecondary peer cooperative learning programs: Annotated bibliography 2019.

Austin P. C. (2011). An introduction to propensity score methods for reducing the effects of confounding in observational studies. Multivariate Behavioral Research, 46(3), 399–424.

Brookhart, M. A., Schneeweiss, S., Rothman, K. J., Glynn, R. J., Avorn, J., & Stürmer, T. (2006). Variable selection for propensity score models. American Journal of Epidemiology, 163(12), 1149–1156.

Carlson, K., Turvold Celotta, D., Curran, E., Marcus, M., & Loe, M. (2016). Assessing the impact of a multi-disciplinary peer-led-team learning program on undergraduate STEM education. Journal of University Teaching & Learning Practice, 13(1), 5.

Chan, J. Y. K. & Bauer, C. F. (2015). Effect of Peer-Led Team Learning (PLTL) on student achievement, attitude, and self-concept in college general chemistry in randomized and quasi experimental designs. Journal of Research in Science Teaching, 52(3), 319–346.

Cheng, D. & Walters, M. (2009). Peer-assisted learning in mathematics: An observational study of student success. Journal of Peer Learning, 2, 23–39.

Frey, R., Fink, A., Cahill, M. J., McDaniel, M. A., & Solomon, E.D. (2018). Peer-led team learning in general chemistry I: Interactions with identity, academic preparation, and a course-based intervention. Journal of Chemical Education, 95(12), 2103–2113.

Gosser, D. K., Cracolice, M. S., Kampmeier, V. R., Strozak, V. S., & Varma-Nelson, P. (2001). Peer-led Team Learning: A Guidebook. Prentice Hall.

Greifer, N. (2021). cobalt: Covariate Balance Tables and Plots. R package version 4.3.1.

Hodara, M. (2013). Improving students’ college math readiness: A review of the evidence on postsecondary interventions and reforms.

Leite, W. (2016). Practical Propensity Score Methods Using R. Sage Publications.

Liu, W., Kuramoto, S. J., & Stuart, E. A. (2013). An introduction to sensitivity analysis for unobserved confounding in nonexperimental prevention research. Prevention Science :The Official Journal of the Society for Prevention Research, 14(6), 570–580.

Lundmark, J., Paradis, J., Kapp, M., Lowe, E., & Tashiro, L. (2017). Development and impact of a training program for undergraduate facilitators of peer-assisted learning. Journal of College Science Teaching, 46(6), 50–54.

Wilson, S. B. & Varma-Nelson, P. (2021). Implementing peer-led team learning and cyber peer-led team learning in an organic chemistry course. Journal of College Science Teaching, 50(3), 44–50.

R Core Team (2019). R: A language and environment for statistical computing. R Foundation for Statistical Computing, Vienna, Austria. URL http://www.R-project.org/

Rose, S. J. (2013). The effectiveness of pre-course and concurrent course interventions on at-risk college physics students’ mechanics performance. (Ph.D. dissertation), University of Illinois at Urbana-Champaign, Urbana-Champaign, IL.

Rosenbaum, P. R. (2002). Covariance adjustment in randomized experiments and observational studies. Statistical Science, 17(3), 286–327.

Sekhon, J. (2011). Multivariate and propensity score matching software with automated balance optimization: the Matching package for R. Journal of Statistical Software, 42(7), 1–52.

Shanbrom, C. (2021). PAL Data. http://webpages.csus.edu/Corey.Shanbrom/PALdata/

Thomas, A. S., Bonner, S. M., Everson, H. T., & Somers, J. (2015). Leveraging the power of peer-led learning: Investigating effects on STEM performance in urban high schools. Educational Research and Evaluation, 21(7-8), 537–557.

U.S. News & World Report. (2021). Campus Ethnic Diversity. www.usnews.com/best-colleges/rankings/regional-universities-west/campus-ethnic-diversity

Wiggins, G. & McTighe, J. (2005). Understanding by Design. Ascd.

Windsor, A., Bargagliotti, A., Best, R., Franceschetti, D., Haddock, J., Ivey, S., & Russomanno, D. (2015). Increasing retention in STEM: Results from a STEM talent expansion program at the University of Memphis. Journal of STEM Education, 16(2), 11–19.

Zhang, Z., Kim, H. J., Lonjon, G., & Zhu, Y. (2019). Balance diagnostics after propensity score matching. Annals of Translational Medicine, 7(1).

